# Towards a species-level phylogeny for Neotropical Myrtaceae: notes on topology and resources for future studies

**DOI:** 10.1101/2023.10.18.562956

**Authors:** The Neotropical Myrtaceae Working Group, Vanessa G. Staggemeier, Bruno Amorim, Mariana Bünger, Itayguara R. Costa, Jair Eustáquio Quintino de Faria, Jonathan Flickinger, Augusto Giaretta de Oliveira, Marcelo T. Kubo, Duane Fernandes Lima, Leidiana Lima dos Santos, Ana Raquel Lourenço, Eve Lucas, Fiorella Fernanda Mazine, José Murillo-A, Marla Ibrahim Uehbe de Oliveira, Carlos Parra-O, Carolyn E. B. Proença, Marcelo Reginato, Priscila Rosa, Matheus Fortes Santos, Aline Stadnik, Amélia Carlos Tuler, Karinne Sampaio Valdemarin, Thais Vasconcelos

## Abstract

**Premise of the study:** Increasingly complete phylogenies underpin studies in systematics, ecology, and evolution. Myrteae (Myrtaceae), with c. 2,500 species, is a key component of the exceptionally diverse Neotropical flora, but given its complicated taxonomy, automated assembling of molecular supermatrices from public databases often lead to unreliable topologies due to poor species identification.

**Methods:** Here, we build a taxonomically verified molecular supermatrix of Neotropical Myrteae by assembling 3,954 published and 959 unpublished sequences from two nuclear and seven plastidial molecular markers. We infer a time calibrated phylogenetic tree that covers 712 species of Myrteae (c. 28% of the total diversity in the clade) and evaluate geographic and taxonomic gaps in sampling.

**Key results:** The tree inferred from the fully concatenated matrix mostly reflects the topology of the plastid dataset and there is a moderate to strong incongruence between trees inferred from nuclear and plastid partitions. Large, species-rich genera are still the poorest sampled within the group. Eastern South America is the best-represented area in proportion to its species diversity, while Western Amazon, Mesoamerica, and the Caribbean are the least represented.

**Conclusions:** We provide a time-calibrated tree that can be more reliably used to address finer-scale eco-evolutionary questions that involve this group in the Neotropics. Gaps to be filled by future studies include improving representation of taxa and areas that remain poorly sampled, investigating causes of conflict between nuclear and plastidial partitions and the role of hybridization and incomplete lineage sorting in relationships that are poorly supported.

## INTRODUCTION

The exceptionally diverse Neotropical flora accounts for more species of flowering plants than tropical Africa and Asia together (Antonelli and Sanmartín, 2011). One of the dominant components in these environments are the myrtles (Myrtaceae), a group of trees and shrubs of outstanding diversity and abundance in both neotropical rainforests (Mori, 1983; Oliveira-Filho and Fontes, 2000; Zipparro et al., 2005; Joly et al., 2012) and savannas (Mendonça et al., 2008; Françoso et al., 2016). Myrtaceae species are dominant in many different kinds of Neotropical ecosystems, such as lowland Atlantic Forest (Staggemeier et al., 2017), rocky outcrop savanna (Santos et al., 2012), high altitude cloud forest and riverine forests (Ibisch et al., 2002). Besides composing a significant part of the biomass that forms these ecosystems, they offer flower resources to pollinators and fleshy-fruits to frugivores and insects (Gressler et al., 2006; Valadão et al., 2019; Martins et al., 2023), which are produced throughout all seasons of the year (Staggemeier et al., 2010, 2015a, 2017). Some species are a promising source of biologically active compounds with medicinal properties such as antimicrobial, antilarval, antioxidant, anti-inflammatory, and cytotoxic (Stefanello et al., 2011; Cascaes et al., 2015). Widespread exploitation of fruits of economic value, such as guava, pitanga, and jaboticaba, make Myrtaceae one of the most used plant families by rural populations showing the high socio-biodiversity value (Souza et al., 2018; Silva et al., 2022). Due to the recent systematic knowledge constructed for the family (Lucas et al., 2007, 2011, 2018, 2019; Mazine et al., 2014, 2016, 2018), Neotropical Myrtaceae have been also elevated from the status of a neglected group to a potential model system in addressing broad questions regarding the dynamics of Neotropical biodiversity (Lucas and Bünger, 2015; Giaretta et al., 2015; Staggemeier et al., 2015b; Vasconcelos et al., 2019a).

In the center of this process is the inference of increasingly robust and complete phylogenetic trees for the tribe Myrteae, the most diverse clade in the family (c. 2,500 species) and the lineage that comprises all new world species (except *Metrosideros stipularis* (Hook. & Arn.) Hook.f. in the tribe Metrosidereae, a species native to Chile). Myrteae entered the phase of molecular systematics in the 2000s with the works of Wilson et al. (2005) and Lucas et al. (2005, 2007), and since then many other studies tackling individual clades and geographic areas (Lucas et al., 2011; Murillo-A et al., 2012; Mazine et al., 2014, Staggemeier et al., 2015b; Bünger et al., 2016; Santos et al., 2016; Wilson et al., 2016; Amorim et al., 2019; Flickinger et al., 2020; Santos et al., 2021; Lima et al., 2021) or reassessing the tribe’s topology in light of a larger species and molecular sampling (e.g. Lucas et al., 2007; Vasconcelos et al., 2017a) have been published (for a complete review see Souza Neto et al., 2022). These studies have provided data and evidence to justify several taxonomic rearrangements (e.g. Proença et al., 2020) and consistent frameworks to shape and test biogeographical, ecological, and evolutionary hypotheses (e.g. Amorim et al., 2019; Bünger et al. 2016; Santos et al., 2017; Staggemeier et al., 2010, 2015b, 2015a; Vasconcelos et al., 2017b, 2018, 2019a).

In spite of these advances, it is not uncommon that larger phylogenetic trees built for the purpose of exploring eco-evolutionary questions (e.g. Seger et al., 2013; Smith and Brown, 2018) contain taxonomic mistakes in this group, for instance by using invalid names or misidentifying samples, potentially leading to topologies that differ strongly from more complete and carefully assembled datasets. The reason for this is that automated methods of tree inference (e.g. Bennett et al., 2018; Smith and Walker, 2019) usually source taxonomic information from public databases of sequences and names that have not been updated at the same pace as the Myrteae taxonomy. Due to poor identification of samples and outdated taxonomy, it is not uncommon that sequences from different vouchers, which are often different species, are assembled together as part of a single tip in tree building. This issue should also impact the interpretation of downstream evolutionary analyses that use these phylogenies (e.g. Antonelli et al., 2018) as well as broad scale compilations of species list (e.g. Ter Steege et al., 2016). A taxonomically verified molecular matrix including all markers that have been most frequently used to infer phylogenies in the group would likely solve these issues and would be an important resource for future finer-scale studies. Much of the molecular data required to produce a satisfactory result in this sense is already produced, but is either scattered throughout several parallel studies or still unpublished. In this sense, coordinating data sharing to build a broad, inclusive, and taxonomically reliable molecular matrix is the most effective way to produce a robust species-level tree for Neotropical Myrtaceae and to make this resource widely available for future studies. Here we retrace the recent developments in the phylogenetics of the Neotropical clade (*sensu* Vasconcelos et al. 2017a) of tribe Myrteae to combine molecular data generated from studies published in the last 15 years plus c. 1,000 unpublished sequences provided by co-authors of this study. We further re-assessed identification of all vouchers to increase taxonomic accuracy and explore the support and consistency of topologies inferred from different molecular markers. We also identify taxonomic and geographical gaps in sampling to be targeted for sequencing and suggest guidelines to improve phylogenetic resolution in future studies. Finally, we provide a densely sampled time-calibrated tree that can be more reliably used to explore future eco-evolutionary venues in the Neotropics. The specific aims of this study are three: (1) to infer a large and, most importantly, taxonomically verified species-level phylogeny of Myrteae; (2) to investigate how to improve species sampling and support for relationships and consistency of topologies inferred from different markers and where to focus resources for future studies; and (3) to provide a calibrated phylogenetic tree with trustworthy species identification that can be used for eco-evolutionary studies that require dated ultrametric trees.

## METHODS

### Taxon sampling

The Neotropical clade of tribe Myrteae, henceforward Neotropical Myrteae, encompasses nine subtribes (*sensu* Lucas et al., 2019): Blepharocalycinae (3 species), Eugeniinae (c. 1,200 species), Luminae (c. 50 species), Myrciinae (c. 750 species), Myrtinae (c. 20 species), Pimentinae (c. 200 species), Pliniinae (c. 120 species), and Ugniinae (12 species). This clade includes c. 2,500 species in total (*sensu* POWO, 2023) of which c. 2,200 are Neotropical. *Eugenia* sect. *Jossinia* (DC.) Nied., a clade of over 200 species within the Eugeniinae, is the largest non-Neotropical radiation within this clade, but it is nested within a clade comprising all Neotropical *Eugenia* L. (Mazine et al., 2016, 2018) and thus is also included as part of the ingroup in our analysis to preserve the monophyly of the group. *Myrtus communis* L. within Myrtinae and *Lophomyrtus* Burret (two species) and *Lenwebbia* N.Snow & Guymer (two species) within Ugniinae are the other non-Neotropical representatives of this group, and were also included as part of the ingroup as they are also nested within clades majoritarily distributed in the Neotropical region (Vasconcelos et al., 2017a). To reconstruct the most species-inclusive phylogenetic hypothesis for Neotropical Myrteae, DNA sequences were retrieved from two sources: (1) by data-mining GenBank and (2) by compiling unpublished sequences generated by the co-authors of this study. Each approach is described below.

### Data-mining GenBank

We searched and downloaded all nucleotide entries for each of the 52 Myrteae genera recognized by Wilson (2011) in February/2023. Given the problematic nature of Myrteae’s taxonomy and to guarantee that all vouchers included in the tree can be traced back to a herbarium for future consultation, several steps of manual curation had to be performed after downloading all sequences available on GenBank. This preliminary list was first filtered to remove sequences not belonging to Myrteae (e.g. sequences from pathogens and parasites that were also occasionally recovered by the search engine). The remaining sequences were classified by subtribe (*sensu* Lucas et al., 2019) and organized in distinct folders in Geneious v. 11 (https://www.geneious.com). Sequences belonging to the subtribe Decasperminae, a non-Neotropical clade sister to all of the remaining extant Myrteae, were also removed except for those of 25 species selected to be part of the outgroup in the phylogenetic inference. Thirty-six species belonging to other Myrtaceae tribes were also searched and included in the dataset for the same purpose. Next, we filtered these results to exclude all but nine molecular markers that showed the greatest coverage among all species: the nuclear regions ITS and ETS and the plastid regions *matK*, *ndhF, psbA-trnH, rpl16, rpl32-trnL, trnL-trnF, trnQ-rps16*. The remaining sequences were renamed as “species_voucher” and reorganized in different folders for individual markers. Sequences for which no voucher information was found or that could not be easily traced back to a herbarium specimen were also excluded in this step. When there were two sequences for the same molecular region and voucher, we kept the one with the longest length. We kept duplicate entries from the same species as long as they belonged to different vouchers. This filtering process resulted in 3,160 sequences recovered from GenBank, in a dataset that is restricted to specimens for which the vouchers are known and can be traced back to a herbarium specimen or living collection.

### Previously unpublished data – extraction and sequencing protocols

Previously unpublished molecular data contributed by co-authors of this study account for 1,024 sequences of the same nine molecular markers targeted in our GenBank search. DNA extraction for most samples was carried out from silica-dried leaf material using the CTAB extraction protocol (Doyle and Doyle, 1987) and left for precipitation under conditions of −18°C in 100% ethanol followed by a purification by equilibrium centrifugation in CsCl-ethidium bromide gradients (1.55g × ml-1). Butanol extraction followed by dialysis were employed to remove the ethidium bromide and caesium chloride. QIAGEN DNeasy Plant Mini Kit (QIAGEN, Germany) was also employed for extracting DNA, following manufacturer’s instructions. The target regions were amplified on the GeneAmp PCR System 9700 (Applied Biosystems, Foster City, California, U.S.A.) and Mastercycler nexus (Eppendorf, Hamburg, Germany). The PCR products were purified using QIAGEN® QIAquick™ PCR Purification Kit according to the manufacturer’s protocol. Sequencing reactions were carried out with the Taq DyeDeoxy™ Terminator Cycle Sequencing Kit (Applied Biosystems, Inc). Sequences were read on an ABI PRISM 3100 Genetic Analyzer (Applied Biosystems, Inc). Primers, PCR and sequencing conditions are detailed in the Supplementary Information 1. All previously unpublished sequences have been deposited in GenBank; DNA samples are deposited in the Royal Botanic Gardens Kew DNA and Tissue Collections.

### Taxonomic verification and alignment

In the next step, all species names and vouchers included in the dataset were circulated among all co-authors to have their taxonomic identities checked and, when required, to update their identifications prior to tree inference analyses. That also comprised vouchers marked as “sp.” (unidentified species) from both GenBank and unpublished data, enabling new identifications to be performed and included. This procedure of manual re-assessment of all data was time-consuming, but is still the most effective way to address the problem of taxonomic inaccuracy that is widespread in Neotropical Myrteae. We then ran alignments for each molecular region separately using the Muscle algorithm (Edgar, 2004) implemented in Geneious, with default settings. Alignments were visually inspected and adjusted for issues with sequence inversions. Sequences that aligned poorly, that were too short (<200 bp) or represented possible contaminations were excluded from the matrix. At the end of this process, the resultant alignment was considered cleaned and used in all the subsequent analyses. Voucher information and GenBank numbers for all sequences utilized will be made available upon publication.

### Tree inference, analyses of support, and time calibration

We first inferred a phylogenetic tree including all the molecular data available in our dataset, comprising a concatenated matrix of all nine molecular markers and all tips (henceforward “the supermatrix tree”). This tree was used to contrast species positions against previously published trees using smaller datasets and to observe general patterns in support and topology among the eight subtribes focused on here. To this end, we used a maximum likelihood (ML) algorithm implemented in RAxML-HPC v.8 on XSEDE v.8.2.12 (Stamatakis, 2006) through CIPRES (Miller et al., 2010), setting it to 1,000 bootstrap replications. We then filtered three subset matrices, complete for all molecular markers: (1) a subset of 53 tips for which all the nine markers were available (henceforward the “all53” set); (2) a subset of 169 tips representing the Myrciinae subtribe (including the large genus *Myrcia* DC.) for which five markers were sampled (i.e. nuclear ITS and the plastid *ndhF*, *psbA-trnH*, *trnL-trnF*, and *trnQ-rps16*) and (3) a subset of 197 tips for which five markers frequently used in studies of the large genus *Eugenia* were sampled (the nuclear ITS and the plastid *psbA-trnH*, *rpl16,* and *rpl32-trnL* and *trnQ-rps16*). Using these subsets, we ran separate ML analyses for each different marker in each subset. We also ran ML analyses for combining partitions from different organelles (“nuclear” and “plastid”) and one analysis for the full-concatenated matrix in each subset (i.e. “full” analysis). All ML analyses were run using the same software and settings as described for the supermatrix tree. Nodes recovered with bootstrap (BS) values of over 75 were considered strongly supported; between 50 and 75 were considered moderately supported and below 50 weakly supported in all cases.

To compare topologies and supports of each marker and combined partition against the full analysis, we used three approaches. First, we extracted all bootstrap values in each tree and compared their median, highlighting which marker inferred trees with highest general support. Then, we estimated the similarity between the tree topology inferred by each marker against the full analysis by extracting a matrix of distances based on the covariance among tips and calculating Mantel statistics between pairs of matrices. The posthoc significance values were estimated following the Pearson method and 1,000 permutations, using functions of the R packages ape (Paradis et al., 2004) and vegan (Oksanen et al., 2007). Values ranged from 0 (topologies are completely different) to 1 (topologies are completely similar). Lastly, we used functions of the R package treespace (Jombart et al., 2017) to simultaneously explore resolution and topology inferred from different markers and partitions, using a sample of 100 trees randomly extracted from the set of bootstrap replications of each ML analysis. This analysis is similar to a multidimensional scaling and visually depicts how different topologies are (i.e. how much they overlap in the multidimensional tree space) and how robust is the support wield from each inference (i.e. the larger the area, the lower the support).

Finally, we produced a time-calibrated tree using the concatenated supermatrix alignment considering two secondary calibration points as well as four fossil constraints based on pollen fossil data. Prior distribution and settings for each calibration point followed “Approach B” in Vasconcelos et al. (2017a). The time calibration was performed in BEAST v.2.7.3 (Bouckaert et al., 2014) through CIPRES. The clock model was selected as “Uncorrelated relaxed clock” and tree prior was selected as “Speciation: birth-death incomplete sampling”. Nucleotide substitution models were set for nuclear and plastid partitions separately. Best substitution models for each partition were selected based on AIC weight using the modelTest function of the R package *phangorn* (Schliep, 2011), resulting in GTR+I+G model for both nuclear and plastid partitions. Due to computational limitations in running BEAST with datasets of this size, we constrained the topology of the calibrated tree to be the same as that recovered in the RAxML analysis. The MCMC was set for 200,000,000 generations, sampling every 10,000 generations. Convergence was assessed using Tracer v.1.7.2 and accepted when the effective sampling size (ESS) was around or above 200 for all parameters. The resultant time-calibrated tree was further pruned to remove outgroups and to leave only one tip per species, resulting in a coverage of 652 species of Neotropical Myrteae (c. 26% of the group). The representative voucher for species with more than one entry in the original tree was selected based on the voucher with most molecular data or which best represents the morphological diagnosis of the species it represents, particularly in cases of species that appear non-monophyletic in the ML tree. All alignments and trees produced are available at https://github.com/tncvasconcelos/myrteae_tree.

### Identifying taxonomical and geographical gaps

To inspect how balanced the final pruned tree is in terms of taxonomic and geographical coverage, we estimated the sampling proportion of species in each subtribe and for each Level-3 Botanical Country based on the categorization of the World Geographical Scheme for Recording Plant Distributions (Brummitt et al., 2001) using data from POWO (2023). This information was used to discuss which clades and geographical areas are particularly over-or undersampled and where to invest resources for future molecular sampling. For discussion on geographical coverage, we focused on the Neotropics since this is the center of diversity in the group with c. 90% of the species and because we noticed some synonym inaccuracies with paleotropical species of *Eugenia* in the POWO dataset.

## RESULTS

### Phylogenetic analyses and time calibration

The final cleaned supermatrix of nine molecular markers contains a total of 4,913 sequences, of which 3,954 (80%) were retrieved from GenBank and 959 (20%) are newly published here. The concatenated alignment has a length of 10,312 base pairs, of which 46% represent missing data; ITS and *psbA-trnH* are the markers most commonly represented throughout the supermatrix. The tree inferred from this supermatrix encompasses 1,030 tips, representing 773 species and 48 genera (including outgroups) and 712 species and all the 34 accepted genera of Neotropical Myrteae (excluding outgroups), c. 28% of the species diversity in the clade (Fig.2,5B). Among the 78 previously unidentified vouchers (i.e. those marked as “sp.” on GenBank), 17 could be newly identified to species level, whereas eight previously misidentified vouchers were newly assigned as “sp.”; 54 vouchers remain unidentified after taxonomic verification. Several species represented by more than one voucher appear polyphyletic, including the widespread *Blepharocalyx salicifolius* (Kunth) O.Berg, *Myrcia amazonica* DC., *Myrcia multiflora* (Lam.) DC., *Myrcia splendens* (Sw.) DC., and *Myrcia guianensis* (Aubl.) DC.

**Figure 1:**
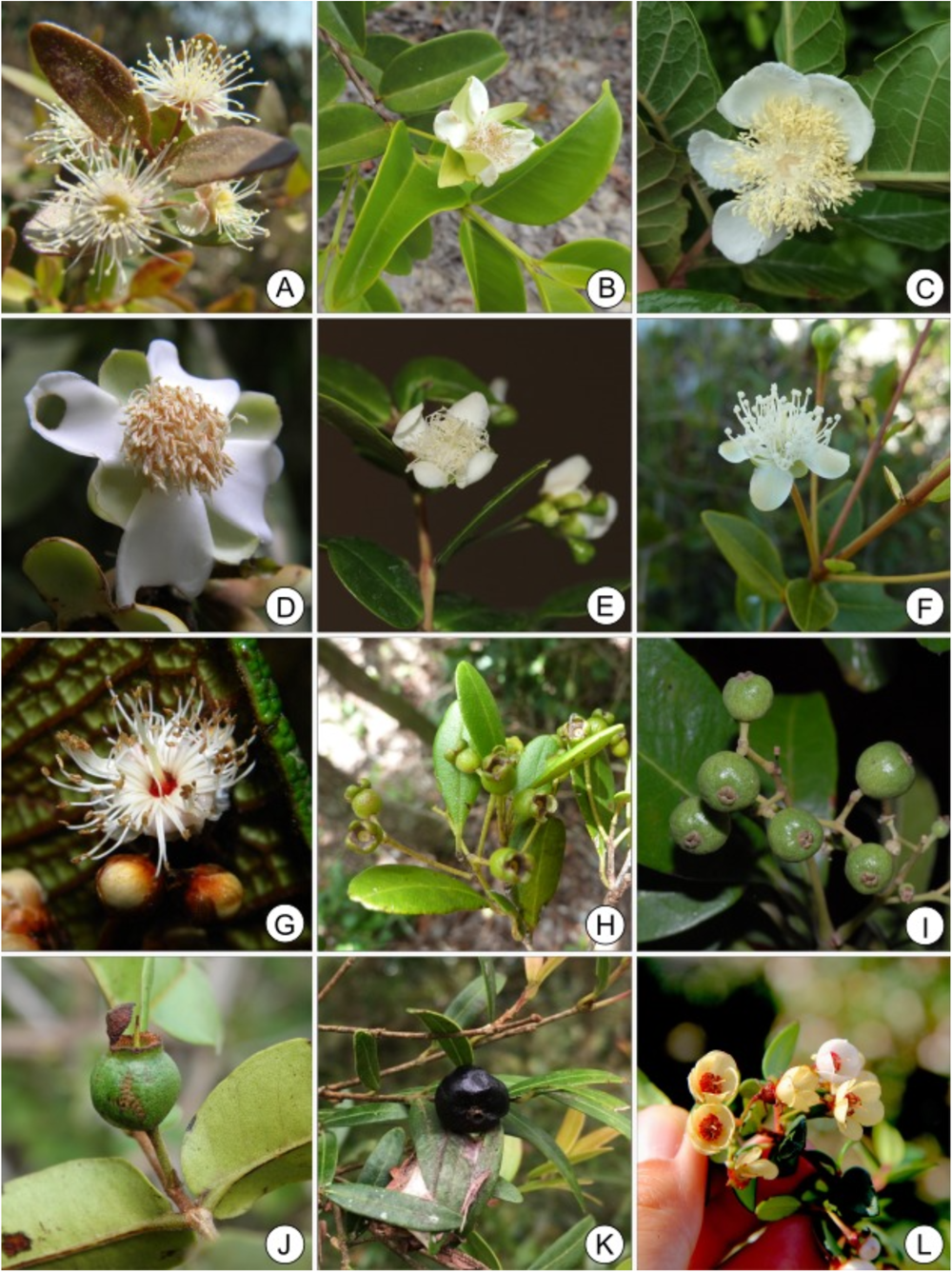
Diversity of flowers and fruits in Myrteae (Myrtaceae). (A) *Blepharocalyx salicifolius* (Kunth) O.Berg (Blepharocalycinae), (B) *Calycolpus legrandii* Mattos (Myrtinae), *Campomanesia schlechtendaliana* (O.Berg) Nied. (Pimentinae), *Eugenia quilombola* B.S.Amorim, M.A.D.Souza & Giaretta (Eugeniinae), *Luma apiculata* (DC.) Burret (Luminae), *Mosiera longipes* (O.Berg) Small (Pimentinae), *Myrcia ruschii* B.S.Amorim (Myrciinae), *Myrcianthes frangrans* (Sw.) McVaugh (Eugeniinae), *Pimenta dioica* (L.) Merr. (Pimentinae), *Psidium brownianum* Mart. ex DC. (Pimentinae), *Siphoneugena reitzii* D.Legrand (Pliniinae), *Ugni candollei* (Barnéoud) O.Berg (Ugniinae). Photo credits: AS (A), MIUO (B, C), BSA (D, G), TV (E, I, L), JF (F, H), AT (J), VGS (K).

**Figure 2:**
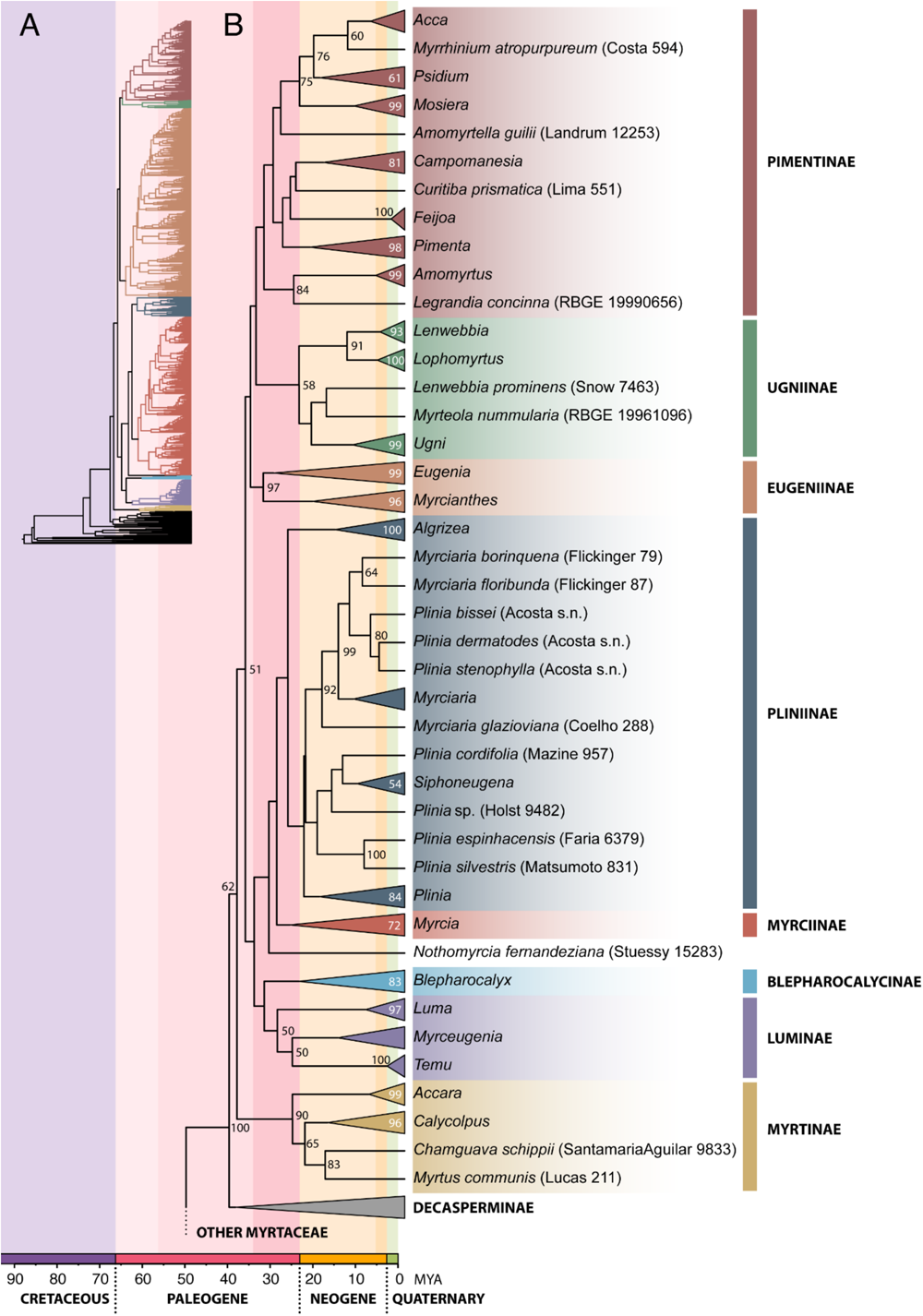
Time calibrated maximum likelihood phylogenetic tree inferred from the concatenated supermatrix of nine markers for Myrteae (Myrtaceae). (A) Shows the full, uncollapsed topology including 1,030 tips. (B) Highlights relationships between the genera and unplaced species comprising the eight predominantly Neotropical subtribes of tribe Myrteae (Myrtaceae). Bootstrap supports <50 are not shown.

Tribe Myrteae is recovered with high support (Fig.2, BS 100) in the ML analysis, as are the recently re-circumscribed subtribes Eugeniinae and Myrtinae (BS 97 and 90 respectively). Myrciinae and the Neotropical clade of Myrteae are recovered with moderate support (BS 72 and 62 respectively) and Pliniinae, Pimentinae, and Luminae all appear as low supported clades (BS < 50). Oldest splits in the phylogeny of Eugeniinae and *Eugenia* appear highly supported (Fig.3) whereas supports tended to be lower for relationships among sections of *Myrcia* in the Myrciinae (Fig.4). Based on estimated ages of the crown groups, the oldest of the Neotropical Myrteae subtribes appears to be Eugeniinae (33.21mya, with a confidence interval (CI) of 30.22-36.12), followed by Pimentinae (33.02mya, 28.92-36.96), Lumiinae (29.89mya, 23.4-35.48), Pliniinae (27.42mya, 23.49-31.48), Myrciinae (26.95mya, 23.39-30.14), Myrtinae (26.33mya, 19.51-32.97), Blepharocalycinae (24.82mya, 17.59-32.58), and finally Ugniinae (24.79mya, 22.12-29.85) (Fig.5A).

**Figure 3:**
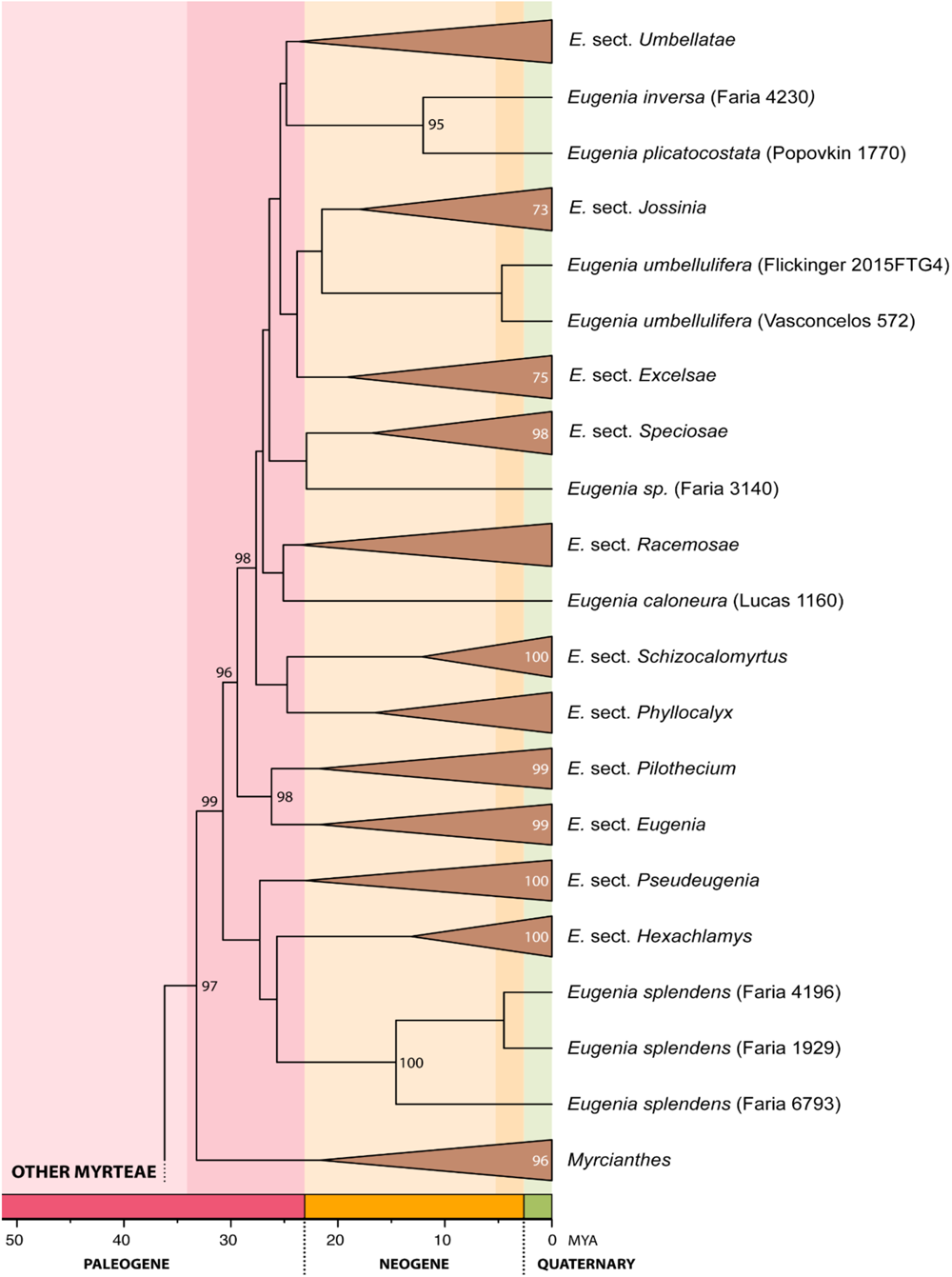
Time calibrated maximum likelihood phylogenetic tree inferred from the concatenated supermatrix of nine markers, highlighting relationships between sections of *Eugenia* and genera within subtribe Eugeniinae. Bootstrap supports <50 are not shown.

**Figure 4:**
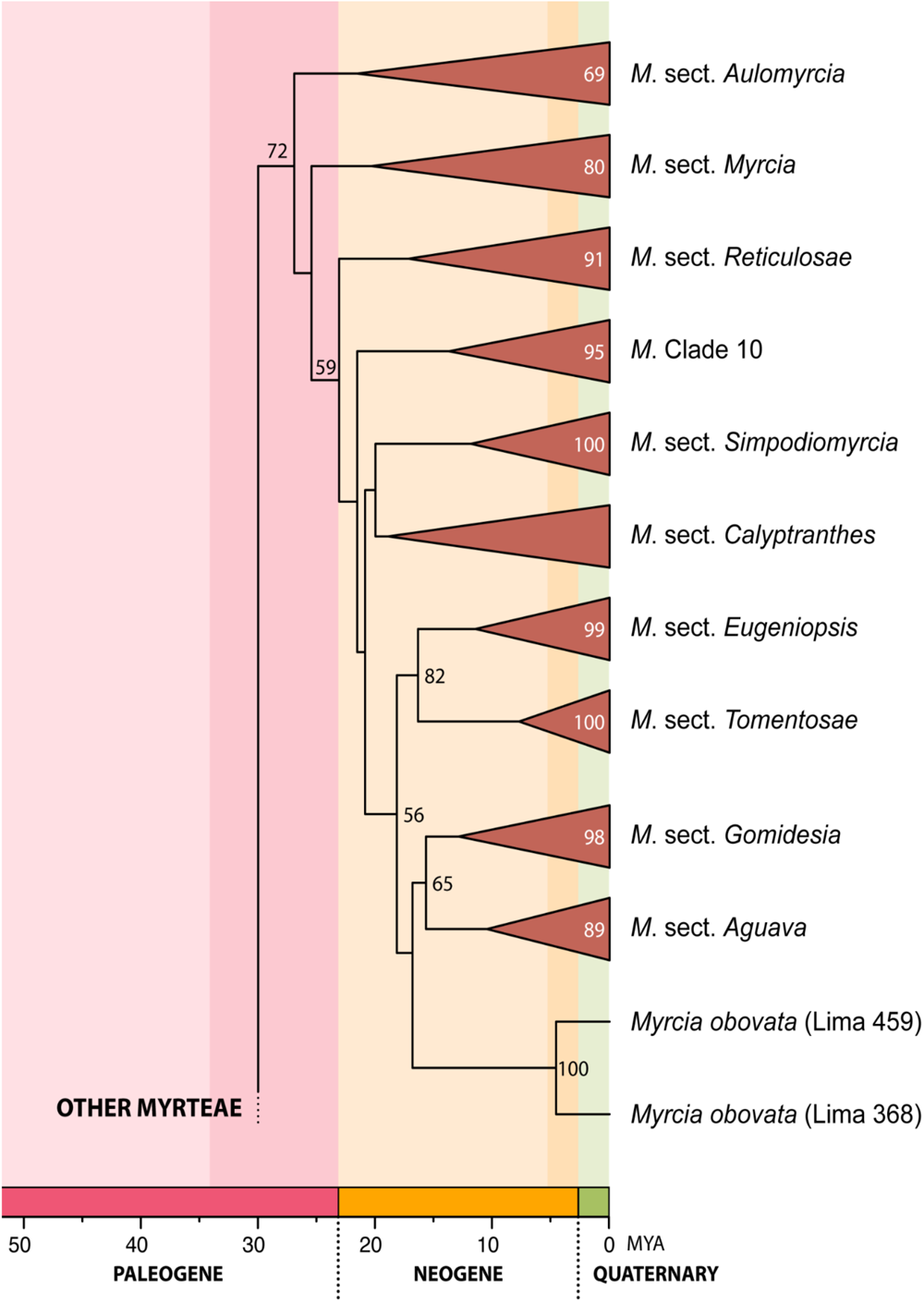
Time calibrated maximum likelihood phylogenetic tree inferred from the concatenated supermatrix of nine markers, highlighting relationships between sections of the mega-diverse genus *Myrcia* (subtribe Myrciinae). Bootstrap supports <50 are not shown.

**Figure 5:**
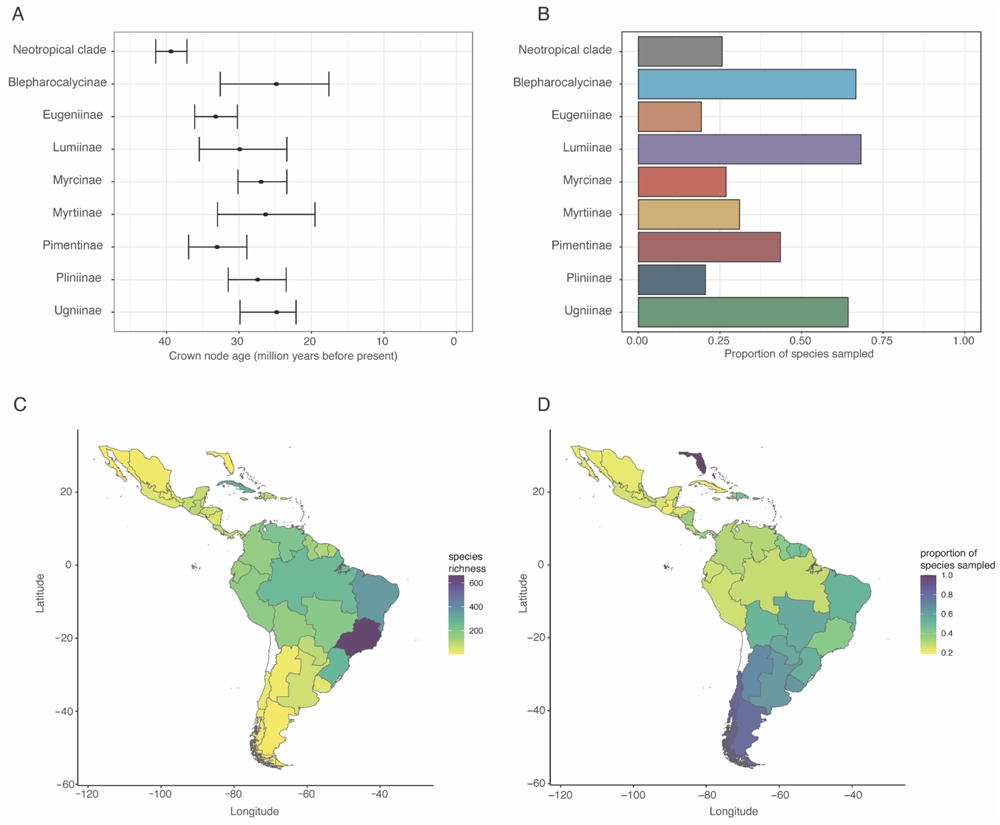
Crown node age and assessment of taxonomic and geographic gap in the phylogenetic sampling. (A) Mean and confidence interval (95%) around age estimates for the crown node of each subtribe; (B) Proportion of species sampled in each subtribe in the tree; (C) Total Myrteae species-richness per level-3 botanical country (TWDG, 2023) based on POWO (2023); (D) Proportion of species sampled in the phylogeny in each level-3 botanical country. No species occurs in Northern Chile, so the area is shown in white.

### Assessment of taxonomic and geographic coverage

Subtribes with the best taxonomic coverage in our supermatrix dataset tend to be the species-poor ones, i.e. those with less than 100 species (Fig.5B). Luminae presents 68% of its c. 50 species sampled, Blepharocalycinae 66% of its three species sampled (all but *Blepharocalyx myriophyllus* (Casar.) P.O.Morais & Sobral) and Ugniinae 64% of its 12 species sampled. The exception is Myrtinae, where only 31% of its c. 20 species have been sampled. The largest subtribes tend to have a lower proportion of their species richness covered, with 16% of the c. 1,200 species of Eugeniinae, 27% of the c. 750 species of Myrciinae, and 21% of the c. 120 Pliniinae sampled in the tree. Pimentinae, a relatively species rich subtribe of c. 200 species, appear relatively well covered, with 44% of its species sampled (Fig. 5C). In terms of proportional sampling across the distribution of Myrteae in the Neotropics, the Amazon basin, Ecuador, Peru, as well as Mesoamerica and some areas in the Caribbean appear particularly poorly sampled in relation to their species richness (Fig. 5C,D).

### Contrasting resolution and topology of each marker

Analyses using the three subsets with complete molecular matrices highlight differences in bootstrap support resulting from analyses of each partition and individual molecular marker and topological conflicts between nuclear and plastid partitions (Fig. 6). In the Myrteae and Myrciinae datasets, full concatenated matrices including all sampled markers consistently present the highest bootstrap support values, followed by the concatenated plastid partitions (Fig. 6A,C). The opposite is observed in the Eugeniinae dataset, where the plastid partition presents higher support than the full concatenated one (see Fig. 6B). Some of the most recurrently used markers in Myrteae phylogenetics, such as *psbA-trnH*, *ndhF, rpl16*, and *trnL-trnF* yielded low support when analyzed in isolation and were highly inconsistent with the full matrix. Surprisingly, the most congruent individual markers in the full analysis are *mat*K and *trnQ-rps16*, two regions less frequently sequenced for the whole tribe. The support yielded by these two markers is even higher than the concatenated nuclear partition in this dataset (Fig. 6A). In general, Mantel test posthoc significance values were congruent to the analyses of support; i.e. phylogenies with lower support also had topologies that were more distinct from the full concatenated one, with a few exceptions (Table 1). Analyses of the phylogenetic landscape shows strong conflict between plastid and nuclear partitions in the full Myrteae dataset, with low overlap between partitions in the treespace (Fig. 6D). This conflict is also observed in the Myrciinae dataset (Fig. 6F), but less so in the Eugeniinae dataset (Fig. 6E) where the plastid partition seems to overlap more closely with the nuclear one. In all datasets, the topology of the full matrix overlaps mainly with the topology inferred from the plastid partition.

**Figure 6:**
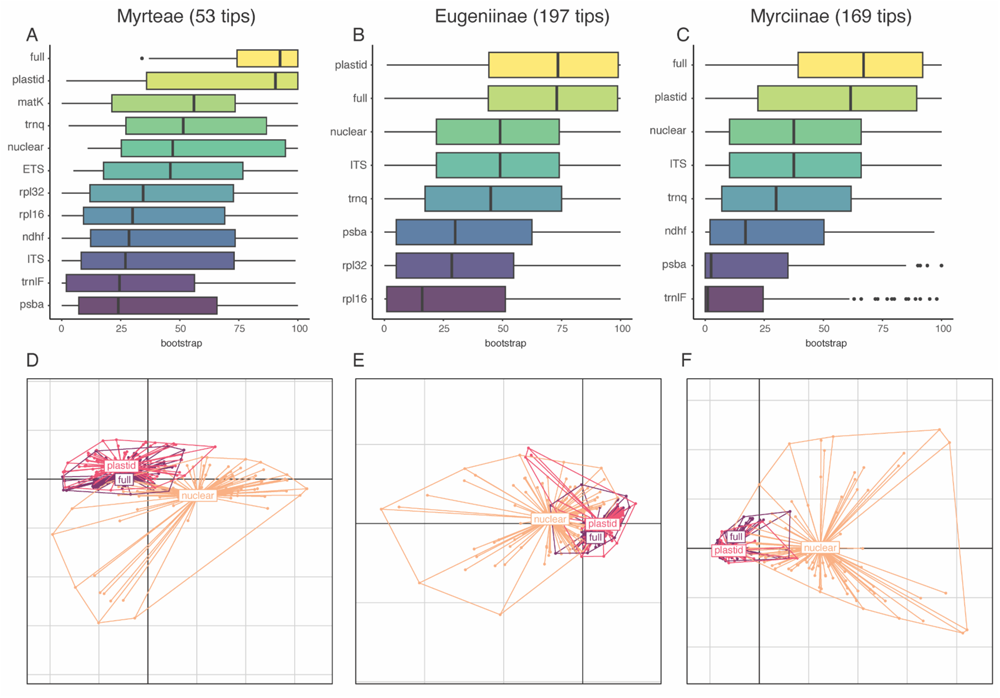
Distribution of bootstrap supports per molecular marker and partition subset and treespaces comparing nuclear and plastid partitions with full datasets. Columns represent subsets for (A,D) Myrteae (53 tips with complete molecular datasets for nine markers); (B,E) Eugeniinae (197 tips with complete molecular dataset for five markers); (C,F) Myrciinae (169 tips with complete molecular dataset for five markers).

**Table 1:** Mantel test posthoc significance values for comparisons between the full concatenated topology and topologies resulting from each individual partition and marker. Values range from 0 (topologies are completely different) to 1 (topologies are completely similar).

|  | nuclear | ETS | ITS | plastid | matK | ndhF | psbA-trnH | rpl16 | rpl32-trnL | trnL-trnF | trnQ-rps16 |
| --- | --- | --- | --- | --- | --- | --- | --- | --- | --- | --- | --- |
| all53 | 0.98 | 0.97 | 0.97 | 1 | 0.96 | 0.95 | 0.91 | 0.88 | 0.93 | 0.91 | 0.94 |
| Eugeniinae | 0.85 |  | 0.85 | 0.99 |  |  | 0.63 | 0.79 | 0.85 |  | 0.83 |
| Myrciinae | 0.82 |  | 0.82 | 0.99 |  | 0.94 | 0.75 |  |  | 0.58 | 0.95 |

## DISCUSSION

### General comments on topology and time calibration

Our inferred phylogeny represents the first Myrtaceae tree to include all the 34 genera of the hyper-diverse Neotropical clade of Myrteae. Most of the resulting suprageneric relationships are congruent with recent phylogenetic analyses performed using different samples within tribe Myrteae (e.g. Vasconcelos et al., 2017a; Maurin et al., 2021), but there are some important differences that must be highlighted when contrasting our results to recently published trees. First and foremost, although our topology recovers all major groups of Myrteae – that is, all subtribes and sections within the large genera *Myrcia* and *Eugenia* (*sensu* Mazine et al., 2016; Lucas et al., 2018) – support across our tree is generally lower than in previous studies that focused on specific groups within the tribe (see discussion below). For practical reasons, our discussion will consider these major groups as clades for now even if support is low and focus on the relationship among and within them. However, given the low support of some relationships at both higher and lower taxonomic levels, it is important to emphasize that our supermatrix tree should not be considered as the final word in terms of phylogenetic relationships within Neotropical Myrtaceae. It is rather a first step towards evaluating sampling gaps and a potential preliminary resource for macroecological and macroevolutionary analyses that need a densely sampled, time calibrated tree.

Though all subtribes (*sensu* Lucas et al., 2019) are recovered as potential clades in our analysis, support varies among them. Specifically, Pliniinae, Pimentinae, and Luminae are all only poorly supported in our supermatrix analysis (BS <50; see all supports in detail for the RAxML tree at Supplementary Information 4). The poor support for Pliniinae might be due to the positioning of the genus *Algrizea* Proença & NicLugh., which has been inconsistently placed as either sister to Myrciinae (*e.g.* in Maurin et al., 2021) or Pliniinae (*e.g.* in Vasconcelos et al., 2017a; Stadnik 2020), depending on the molecular dataset used in the inference (Lucas et al., 2007). Furthermore, the support for Pliniinae as sister to Myrciinae, although recovered, is only poorly supported here (BS <50). Genera included in Pimentinae have been previously found to be paraphyletic in other tribe-level analyses (*e.g.* Vasconcelos et al., 2017a, which divided them into informal *Pimenta* and *Psidium* groups) and the group has also been poorly supported in phylogenomic analyses of the order Myrtales (Maurin et al., 2021). In the tree inferred here, relationships within *Psidium* L. were reasonably consistent with those recovered in the recent phylogeny of the genus (Proença et al., 2022), though with lower support. The monophyly of the four sections of *Psidium* was confirmed, although the relative position of *Psidium* sect. *Obversifolia* O.Berg differed from that of their tree. *Pimenta pseudocaryophyllus* (Gomes) Landrum is recovered together with the rest of *Pimenta*, contrary to Maurin et al. (2021). This is an important difference because the placement of *P. pseudocaryophyllus* in the Myrtinae subtribe in the latter study indicated the possible resurrection of the monotypic genus *Pseudocaryophyllus* O.Berg. Pimentiinae is found along the whole of the Myrteae geographic distribution in the Neotropics and includes genera with wide variation in embryo (Landrum and Stevenson, 1986), floral (Vasconcelos et al., 2015; 2019b) and fruit traits (Pittarelli et al., 2021), indicating that it might in fact constitute more than one natural lineage. The issue with the monophyly of Pimentinae has been discussed when the subtribe was first proposed (i.e. Lucas et al., 2019) and it definitely deserves to be the focus of future studies focused on this clade, especially given the economic importance of members of the group (e.g. the all-spice *Pimenta dioica* (L.) Merr., the guava *Psidium guajava* L., and the aggressive invasive *Psidium cattleyanum* Sabine).

Some general comments are also required regarding the topologies of the super diverse Eugeniinae and Myrciinae subtribes (Figs. 3, 4). Due to the size and sparse nature of the molecular matrix, relationships among sections within the large *Eugenia* and *Myrcia* also presented generally lower support in the full dataset than those recovered by recently published studies focused on fewer species and using more complete molecular datasets. The relationships among *Myrcia* sections, for instance, received higher support in the supermatrix analysis of Amorim et al. (2019). In our analysis, *Myrcia* sect. *Aulomyrcia* (O.Berg) Griseb. appears as a clade with low support (BS 69) sister to the rest of *Myrcia*, followed by *Myrcia* sect. *Myrcia* (BS 80) as sister to the remaining sections. *Myrcia* sect. *Aulomyrcia* has not been consistently recovered as a clade in previous analyses (e.g. Santos et al., 2017) and the reason might be linked to partition conflict (see below). Both *M*. sect. *Aulomyrcia* and *M*. sect. *Myrcia* are the most geographically widespread sections of *Myrcia*, occurring across the whole Neotropical region, whereas the diversity of species in other sections is mainly concentrated in the Cerrado and Atlantic Forest domains of Eastern Brazil (except perhaps for *M*. sect. *Calyptranthes* (Sw.) A.R.Lourenço & E.Lucas which is also diverse in the Caribbean region). A better understanding of the relationships in these earliest splits and across the backbone of *Myrcia* is necessary for untangling the biogeographical history of the group in the Neotropics. Among other sections, *M*. sect. *Calyptranthes* is recovered with low support (BS <50) and placed as sister to *M*. sect. *Sympodiomyrcia* M.F.Santos & E.Lucas (BS 100) as in previous studies (e.g. Santos et al., 2016; 2017). Although this relationship is also poorly supported, these two sections share some morphological characteristics, such as sympodial branching, cataphylls, and deciduous calyx lobes (Santos et al., 2016). *Myrcia* “Clade 10” (BS 95) (Amorim et al., 2019; Lima et al., 2021) is recovered with high support. *Myrcia* sect. *Gomidesia* (O.Berg) B.S.Amorim & E.Lucas (BS 98) is recovered with high support, and so is *Myrcia* sect. *Aguava* (Raf.) D.F.Lima & E.Lucas (BS 89), as long as *M. obovata* (O.Berg) Nied. is not included as part of the latter. These two sections are recovered as forming a clade, as supported by Lucas et al. (2011) and Amorim et al. (2019). *Myrcia obovata* appears as sister to this clade in our tree, contradicting previous molecular analyses and morphological characteristics that place this species within *M.* sect. *Aguava* (Lima et al., 2021). *Myrcia* sect. *Eugeniopsis* (O.Berg) M.F.Santos & E.Lucas (BS 99) is recovered as sister to *M*. sect. *Tomentosae* E.Lucas & D.F.Lima (BS 100) with high support (BS 82), consistent with previous studies (Amorim et al., 2019; Santos et al., 2016, 2017). Relationships within each section are mostly consistent with previous studies focused on specific groups (e.g. Wilson et al., 2016; Santos et al., 2017; Amorim et al., 2019; Santos et al., 2021, Lima et al., 2021), with the exception of *M*. sect. *Aulomyrcia*, as stated above.

The reconstructed phylogenetic hypothesis of Eugeniinae mostly confirms the general topology proposed by Mazine et al. (2014, 2018), Vasconcelos et al. (2017a) and Giaretta et al. (2022). *Eugenia* sect. *Jossinia* (BS 73), the largest non-neotropical clade in our sample, is geographically structured. The relationships in this clade suggest a westward route of round-the-world colonization, with a clade composed of Southeast Asia and New Caledonia species sister to species distributed in Mauritius, India, Madagascar and Africa (Supplementary Information 4). Disjunct geographical patterns in this and other Myrteae groups are likely a result of long-distance dispersal events or dispersal through Antarctica before the Miocene, when the ice sheet that covers that continent was formed (Vasconcelos et al., 2017a; Estrella et al., 2019). Because these relationships are poorly supported, further studies that improve the resolution of the relationships within this group are necessary to confirm potential biogeographical routes.

In terms of time estimates, divergence ages among subtribes are generally in accordance with previous inferences for this group that used a similar fossil dataset for time calibration (*e.g.* Thornhill et al., 2015; Vasconcelos et al., 2017a). It is interesting to note that, when confidence intervals are considered, all subtribes have very similar age, and have likely originated in the Oligocene (33-23 mya). It is worth noting that our calibration approach used only pollen fossil data, given that their systematic placement has been recently thoroughly reviewed (Thornhill et al., 2012). However, a promising way forward in improving our current understanding of time estimates is by reviewing the macrofossil record for Myrteae as well. Several macrofossils assigned to Myrtaceae have been recovered from the Cretaceous of Southern South America, with some being placed within tribe Myrteae (*e.g.* Poole et al., 2001; Ragonese, 1980); including them in future calibrations analyses may push back the crown age of Myrteae (*e.g.* Murillo-A et al., 2016). Though that means that time divergences may change once more fossil data and better time divergence models are incorporated in the analyses, we believe the current time calibrated, taxonomically verified tree can be preliminarily used as a resource for novel studies on ecology and evolution until further data is available. Other ways forward in improving the tree are discussed below.

### Next steps on phylogenetics of Neotropical Myrteae: tackling lack of resolution and uneven species sampling

The low bootstrap support in our supermatrix tree likely results from using a sparse and patchy molecular matrix, which contains several gaps that occasionally reduce branch support. Evidence for this is that the mean bootstrap support increases when we analyze subsets with complete molecular sampling for Myrteae, Myrciinae, and Eugeniinae (i.e. Fig. 5a,c,e). Analysis of closely related lineages in a subset is likely to incorporate less genealogical discordance and avoid genuine phylogenetic signal being blurred by stochastic processes such as gene duplication and loss, hybridization, and incomplete lineage sorting (see Smith et al., 2015; Molloy and Warnow, 2018). In *Eugenia*, a more data rich sample obtained from targeted sequencing recognized that conflicting signal are involved in the low support recurrently recovered along the backbone (Giaretta et al., 2022), which suggests that incomplete lineage sorting or other stochastic processes may play a role in Myrteae phylogenetics. Rapid radiation and recent divergence have been implicated in phylogenetic incongruence and incomplete lineage sorting (Cai et al., 2021; Meleshko et al., 2021) and further studies will be able to determine if this is also the case in Myrteae.

Another potential reason for the lower resolution in the concatenated supermatrix is the conflict between plastid and nuclear partitions. When comparing topologies resulting from analyses of different markers we observe that topologies inferred from nuclear and plastid datasets are strongly incongruent in the full dataset (Figure 5b,d,f). Diverging topologies between plastid and nuclear datasets are expected given that evolutionary rates in nuclear partitions can be five times faster than chloroplast ones in angiosperms (Drouin et al., 2008). However, the incongruence observed between nuclear and full datasets, as well as the overlap between plastid and full topologies in our treespace analysis, may be due to the plastid partition being more data-rich (i.e. longer) than the nuclear one. These analyses show that our full concatenated topology is probably being driven mostly by the plastid partition, where most of the data comes from. That could explain some small inconsistencies between this and the Maurin et al. (2021) genus-level topology using the Angiosperms-353 probe set, which is a nuclear-only dataset (Johnson et al., 2019). Examples of these inconsistencies include the placement of *Algrizea* and *Pimenta pseudocaryophyllus* as discussed above and support for Ugniinae as a clade (recovered in ours but not in Maurin et al., 2021). However, given that most higher level relationships are congruent between our supermatrix and Angiosperms-353 topologies (Maurin et al., 2021; Giaretta et al., 2022) and that Angiosperms-353 topologies generally result in higher node support, we believe that using the Angiosperms-353 in a broader Myrteae sample is a promising way forward to improve our current understanding of relationships in the group. This method will also open the opportunity of contrasting complementary datasets to best assess deep and shallow phylogenetic levels as well as perform relevant genomic analyses to recognize if other processes than genuine conflict signals are taking place. For instance, Giaretta et al. (2022) yielded several infrageneric level topologies of *Eugenia* contrasting different datasets (exons, introns and off-target plastid) and reconstructions methods based on the Angiosperms-353. Even though their general topology agrees with the previous phylogenetic reconstructions (e.g. Mazine et al., 2014, 2018), some inconsistencies were better understood, such as the relationships between groups within *E.* sect. *Umbellatae* O.Berg.

Another important way forward to a better understanding of relationships in the tribe is to increase species sampling, particularly in clades and areas that are currently undersampled. For instance, we show that the largest subtribe in Myrteae, Eugeniinae, remains with less than a fifth of its diversity with trustworthy sequences available for integrative phylogenetic analyses. Sampling is also uneven within *Eugenia*, which makes up most of Eugeniinae with over 1,200 species: the gigantic *Eugenia* sect. *Umbellatae* (c. 500 species) has only c. 12% of its species-richness sampled in our phylogeny. The second largest section, *Eugenia* sect. *Jossinia* is also poorly represented, with only 37 out of the estimated 200 species sampled in our tree. There is a bias in geographical collecting effort as well. Some areas of Southern and Eastern South America are better represented than the Amazon basin and Mesoamerica. Furthermore, although Eastern Brazil appears relatively well sampled (Fig.5D), it is likely that many of the unsampled species in the tree occur in this region, given the exceptional species richness of the group in these areas (Fig.5C). Increased sampling from these areas will also help shed light on species delimitation of widespread species. Some *Myrcia* species (*e.g.*, *Myrcia amazonica*, *M. guianensis*, *M. multiflora*, and *Myrcia splendens*) are not monophyletic on the full analysis and in the Myrciinae dataset. These species are widely distributed across several ecosystems in the Caribbean, Mesoamerica, and South America, and most of them exhibit high morphological variability. Further analyses are needed for understanding if such constitute separate lineages or not, and if so if they could be splitted into two or more species, according to clades obtained in the phylogeny. Given some reported difficulties in extracting sequences from herbarium material even using target sequencing approaches (e.g. Brewer et al., 2019), more field collections will probably be required for such endeavors, especially focusing on groups that lack sampling and in poorly collected areas. New collaborative efforts with botanists in underrepresented regions will represent a key step forward towards this goal.

### Conclusions

Here we assembled a densely sampled molecular supermatrix for the Neotropical clade of tribe Myrteae, Myrtaceae. We showed that reassessing the identification of vouchers used in other studies improved identification accuracy of currently available data, which is of extreme importance to diminish biases in analyses using taxonomically complex groups such as this. The careful taxonomic validation of datasets of this size can be time-consuming, but it still is the best approach for assembling trustworthy phylogenies of taxonomically complicated groups such as Myrteae. Working toward an extended specimen network could facilitate taxonomic updates of other specimen-derived data and research products (Lendemer et al., 2020).

We also provide c. 1,000 new sequences, representing one fifth of the currently available molecular information in the group, which can now be reliably used for other future ecological and evolutionary inferences in this important tropical group. This process resulted in a broadly sampled time calibrated tree that is expert-verified by taxonomists and available for use by ecologists and evolutionary biologists to test hypotheses in their fields, avoiding data of dubious identifications. Gaps for future studies are highlighted; addressing these will involve resolving the incongruence between plastid and nuclear partitions, including a broader sample of species from mega-diverse groups and poorly sampled areas as well as overcoming noise from stochastic processes such as hybridization, introgression or incomplete lineage sorting to better accommodate gene tree heterogeneity. These future endeavors may provide resolution for some persistent taxonomic issues highlighted in the discussion. Results presented here guide future studies to clades that are most urgently in need of sampling. Given the difficulty in accessing suitable samples for sequencing, compiling large phylogenies of tropical groups is a challenge. However, these phylogenies represent a window for greater understanding of highly diverse, taxonomically complex groups from tropical ecosystems where these lineages are often most species rich.

## ACKNOWLEDGEMENTS

VGS was funded by FAPESP (16/02312-8), CNPq (473468/2010-7 and 236687/2012-3), CAPES PhD fellowship, Bentham-Moxon Trust, and Serrapilheira (R-2111-39858). BSA was funded by CAPES (88882.315044/2019-01). DFL is funded by CAPES (88887.371829/2019-00). CEBP, FFM, and MR are funded by CNPq (305105/2022-1, 314593/2020-9, and 302490/2022-1 respectively). I.R.C., V.G.S., and C.E.B.P. also received funding from CNPq through the project “Estudos filogenéticos e macroecológicos voltados à conservação do gênero *Psidium* L. (Myrtaceae Juss. – 2011-2015, REFLORA Program) and I.R.C. through KLARF (Kew Latin American Research Fellowship). Authors acknowledge staff from herbaria ASE, HUEFS, K, SPF, UB, and collection permits from SISBIO (21120-1 to MIUO, 41156-6 to DFL); Laszlo Csiba, Dion Devey, Jim Clarkson, Laura Martinez, J. Floriano Pastore and Maria José G. de Andrade provided laboratorial support; Rob Smissen and Marcos Sobral read and provided comments that greatly improved the manuscript.

## Notes

https://github.com/tncvasconcelos/myrteae_tree

### Competing Interest Statement

The authors have declared no competing interest.

