## Supplementary Information 1 for "Towards a species-level phylogeny for Neotropical Myrtaceae: notes on topology and resources for future studies"

**Supplementary Information 1**: Primers, PCR and sequencing conditions for molecular markers used in the phylogenetic inferences.

| **Regions** | **Primers** | **Primer Sequence (5’-3’)** | **Source** |
| --- | --- | --- | --- |
| ETS | ETS-9 bp | CATGGGCGTGTGAGTGGTGA | Wright et al. (2001) |
|  | ETS-18S | GAGCCATTCGCAGTTTCACAG | Wright et al. (2001) |
|  | ETSMyrt-F | CTCCGTGCTGGTGCATCGAACTGC | Lucas et al. (2007) |
| ITS | ITS92 | AAGGTTTCCGTAGGTGAA | Desfeaux et al. (1996) |
|  | ITS75 | TATGCTTAAACTCCACGGG | Desfeaux et al. (1996) |
|  | ITS18_F | GTCCACTGAACCTTATCATTTAGAGG | Beyra-Matos (1999) |
|  | ITS26_R | GCCGTTACTAAGGGAATCCTTGTTAG | Kass & Wink (1997) |
|  | P1L | CTGTAGGTGAACCTGCGGAAGGATC | Crisp et al. (1999) |
|  | P2R | CTTTTCCTCCGCTTATTGATA | Crisp et al. (1999) |
|  | ITS s4 | TAGCCCCGCCTGACCTGAGG | Kass & Wink (1997) |
|  | ITS s3 | AACCTGCGGAAGGATCATTG | Kass & Wink (1997) |
|  | ITS_SE17_F | ACGAATTCATGGTCCGGTGAAGTGTTCG | Douzery et al. (1999) |
|  | ITS_SE26_R | TAGAATTCCCCGGTTCGCTCGCCGTTAC | Douzery et al. (1999) |
|  | ITS2F | ATGCGATACTTGGTGTGAAT | Schultzj et al. (2007) |
|  | ITS2R | GACGCTTCTCCAGACTACAAT | Schultzj et al. (2007) |
|  | ITSa F | CCTTATCATTTAGAGGAAGGAG | Schultzj et al. (2005) |
|  | ITS-5 | GGAAGTAAAAGTCGTAACAAGG | White et al. (1990) |
|  | ITS-4 | TCCTCCGCTTATTGATATGC | White et al. (1990) |
|  | ITS-3 | GCATCGATGAAGAACGCAGC | White et al. (1990) |
|  | ITS-2 | GCTGCGTTCTTCATCGATGC | White et al. (1990) |
|  | ITS-1 | TCCGTAGGTGAACCTGCGG | White et al. (1990) |
|  | ITSAB101-F | CGAATTCATGGTCCGGTGAAGTGTTCG | Sun et al. (1994) |
|  | ITSAB102-R | GAATTCCCCGGTTCGCTCGCCGTTAC | Sun et al. (1994) |
| *psbA – trnH* | psbA-F | GTTATGCATGAACGTAATGCTC | Sang et al. (1997) |
|  | trnHR | CGCGCATGGTGGATTCACAAATC | Sang et al. (1997) |
|  | trnH2 | CGCGCATGGTGGATTCACAATCC | Tate & Simpson (2003) |
|  | psbA | CGAAGCTCCATCTACAAATGG | Hamilton (1999) |
|  | trnH (GUG) | ACTGCCTTGATCCACTTGGC | Hamilton (1999) |
|  | MYpsb A 1 | TTTTGATTGCAAAATAAAGGAGCAA | Staggemeier et al. (2015) |
|  | rpl2 | GATAATTTGATTCTTCGTCGCC | Goulding et al. (1996) |
|  | eucpsbA | GGAGCAATAACCAACACTCTTG | Freeman et al. (2001) |
|  | trnHf_05 | CGCGCATGGTGGATTCACAATCC | Tate & Simpson (2003) |
|  | 3f | GTTATGCATGAACGTAATGCTC | Cuénoud et al. (2002) |
|  | f | CGCGCATGGTGGATTCACAATCC | Cuénoud et al. (2002) |
|  | trnH-psbA F | TGATCCACTTGGCTACATCCGCC | Shinozaki et al. (1986) |
|  | trnH-psbA R | GCTAACCTTGGTATGGAAGT | Speilmann et al. (1983) |
|  | F | CGAAGCTCCATCTACAAATGG | Kress & Ericson (2007) |
|  | F | ACTGCCTTGATCCCACTTGGC | Kress & Ericson (2007) |
| *rpl16* | Rpl16-F71 | GCTATGCTTAGTGTGTGACTCGTTG | Jordan et al*.* (1996) |
|  | Rpl16-R1516 | CCCTTCATTCTTCCTCTATGTTG | Small et al. (1998) |
| *rpl32 - trnL* | rpl32-F | CAGTTCCAAAAAAAACGTACTTC | Shaw et al. (2007) |
|  | MytrnL(UAG) | CGTTTTCGTAGTTTATGCTCTCCT | Faria et al. (2014) |
|  | Myrpl32-F | ACAAGATGTTCAGTTCAGGCCA | Faria et al. (2014) |
|  | trnL(UAG) | CTGCTTCCTAAGAGCAGCGT | Shaw et al. (2007) |
| *trnQ – rps16* | trnQ (UUG) | GCGTGGCCAAGYGGTAAGGC | Shaw et al. (2007) |
|  | MytrnQ-R | AGTTGATGTAAAGGAAGATTTAGACTC | Murillo-A et al. (2012) |
|  | Myrps16-F | GCGTAAAAWGAGGAAATGCTTAATG | Murillo-A et al. (2012) |
|  | rps16x1 | GTTGCTTTYTACCACATCGTTT | Shaw et al. (2007) |
| matK | matK390-F | CGATCCTTTCATGCATT | Johnson & Soltis (1994) |
|  | matK1326-R | GTATTAGGGCATCCCATT | Johnson & Soltis (1994) |
|  | matK 700F | CAATCTTCTCACTTACGATCAACATC | Gruenstaeudl et al. (2009) |
|  | matK1710R | GCTTGCATTTTTCATTGCACACG | Samuel et al. (2005) |
|  | trnK-R3 | CGGGGCTCGAACCCGGA | Wicke & Quandt (2009) |
|  | trnK2R | CCCGGAACTAGTCGGATGG | Steele &Vilgalys (1994) |
|  | matK 909 F | GGGGTTGCTAACTACACGG | Lam et al. (2002) |
|  | matK 2518 R | TCTGTTGATACATTCGAGTA | Gadek et al. (1996) |
|  | matK 2516 F | TATGCACTTGCTCATGATCA | Gadek et al. (1996) |
|  | matK 2519 R | TTTACGAGCCAAAGTTTTAA | Gadek et al. (1996) |
|  | matK 2521 F | TTCACATTTAGATTATGTG | Gadek et al. (1996) |
|  | matK 756R | ACATATATGAGAATTATATAGG | O’Brien et al. (2000) |
|  | matK 3F_Kim f | CGTACAGTACTTTTGTGTTTACGAG | Ki-Joong Kim, Unpublished |
|  | matK 1R_Kim r | ACCCAGTCCATCTGGAAATCTTGGTTC | Ki-Joong Kim, Unpublished |
|  | matK 5R | GTTCTAGCACAAGAAAGTCG | Ford et al. (2009) |
|  | matK 2.1F | CCTATCCATCTGGAAATCTTAG | Ford et al. (2009) |
|  | matK 2.1a | ATCCATCTGGAAATCTTAGTTC | Cowan et al. (2006) |
|  | matK 3.2r | CTTCCTCTGTAAAGAATTC | Cowan et al. (2006) |
|  | matK 390f | CGATCTATTCATTCAATATTTC | Cuenoud et al. (2002) |
|  | matK 1326r | TCTAGCACACGAAAGTCGAAGT | Cuenoud et al. (2002) |
|  | trnK685F | GTATCGCACTATGTATCATTTGA | Wojeiechowski et al. (2004) |
|  | trnK2R | GTTCTAGCACAAGAAGTCG | Wojeiechowski et al. (2004) |
|  | Kew XF | TAATTTACGATCAATTCATTC | Soltis et al. (2001) |
|  | matKpkF1 | TTTCTGATGAACAARTGGAA | Fazekas et al. (2008) |
|  | matKpkR1 | CGTATCGTGCTTTTRTGYTT | Fazekas et al. (2008) |
| ndhF | 1252-F | GATGAAATTMTTAATGATAGTTGGT | Cook et al. (2008) |
|  | 2063-R | CATTTGGAATTCCATCAATTA | Biffin et al. (2006) |
|  | 748f | CAGTTGCTAAATCGGCACAATT | Biffin et al. (2006) |
|  | 1318r | CGAAACATATAAAATGCRGTTAATCC | Olmstead and Sweere (1994) |
|  | mel-r1 | ATATTGCATAAAAAGCATCTAT | Cook et al. (2008) |
|  | ndhF | GAAAGGTATKATCCAYGMATATT | Shaw et al. (2007) |
|  |  | TCTTTTCTTTTAAGTCTATTTCTTCT | Flickinger et al. (2020) |
|  |  | TCATTAACTAATTCATGGTAGAAC | Flickinger et al. (2020) |
| trnL-trnF | c B49317 | CGAAATCGGTAGACGCTACG | Taberlet et al. (1991) |
|  | d A49855 | GGGGATAGAGGGACTTGAAC | Taberlet et al. (1991) |
|  | e B49873 | GGTTCAAGTCCCTCTATCCC | Taberlet et al. (1991) |
|  | f A50272 | CGAAATCGGTAGACGCTACG | Taberlet et al. (1991) |

| **Pcr conditions** | **ITS** | **ETS** | **psbA-trnH** | **rpl16** | **rpl32-trnL** | **trnQ-rps16** | **ndhF** | **trnL-trnF** | **matK** |
| --- | --- | --- | --- | --- | --- | --- | --- | --- | --- |
| Initial Denaturation  and Taq activation | 94°C (2 min.)/  94°C (5 min.) | 94°C (4 min.) | 80°C (5 min.)/  94°C (4 min.)/  94°C (3 min.)/  94°C (5 min.)/  98°C (45 sec.) | 80°C (5 min.)/  94°C (5 min.) | 80°C (5 min.)/  94°C (5 min.)/ | 80°C (5 min.)/  94°C (5 min.)/ | 80°C (5 min.)/  94°C (4 min.)/ | 95°C (3 min.)  80°C (5 min.)/ | 80°C (5 min.)/  94ºC (30sec.)/  94ºC (1min.)/  94°C (3 min.)/ 94°C (5 min.)/  95°C (2 min.)/ |
| Denaturation** | 91°C (1 min.)/ 94°C (1 min.)/  94°C (20 sec.) | 94°C (1 min.) | 95°C (1 min.)/  94°C (30 sec.)/  94°C (1 min.)/  94°C (45 sec.)/  98°C (10 sec.) | 95°C (1 min.)/  94°C (1 min.) | 95°C (1 min.)/  94°C (1 min.) | 95°C (1 min.)/  94°C (1 min.) | 95°C (1 min.)/  94°C (1 min.) | 95°C (1 min.)  94°C (1 min.) | 95°C (1 min.)/  94ºC (1min.)/  94°C (45 sec.)/  94°C (20 sec.)  95°C (30 sec.) |
| Annealing** | 50°C (1 min.)/  52°C (1 min.)/  48°C (1 min.)/  50°C (30 sec.) | 50°C (1 min.) | 48°C (1 min.)/  52°C (30 sec.)/  56°C (40 sec.)  55°C (45 sec.)/  64°C (30 sec.) | 50°C (1 min.)/  48°C (1 min.)/ | 50°C (1 min.)/  48°C (1 min.)/ | 50°C (1 min.)/  48°C (1 min.)/ | 50°C (1 min.)/ | 50°C (1 min.) | 50°C (1 min.)/  46ºC (40 sec.)/ 48ºC (40 sec.)/  55°C (45 sec.)/  50°C (30 sec.)/  57°C (1 min.)/  52°C (20 sec.) |
| Extension** | 72°C (1:30 min.)  72°C (1 min.)  72°C (45 sec.) | 72°C (1 min.) | 65°C (5 min.)  72°C (1 min.)/  72°C (2:30 min.)/  72°C (2 min.)/  72°C (40 sec.) | 65°C (4 min.)*/  72°C (1 min.) | 65°C (4 min.)*/  72°C (1 min.) | 65°C (4 min.)*/  72°C (1 min.) | 65°C (5 min.)/  55°C (1 min.)  72°C (2 min.) | 65°C (1:30 min.)  72°C (2 min.) | 65ºC (5 min.)/  72ºC (40 sec.)/  72ºC (1min.)  72°C (2 min.)  72°C (45 sec.)  72°C (2:30 min.)/  72°C (50 sec.) |
| Final Extension | 72°C (7 min,)  72°C (4 min,) | 72°C (7 min.) | 65°C (4 min.)/  72°C (10 min.)/  72ºC (3min.)/  72ºC (7min.) | 65°C (4min.)/  72ºC (7min.) | 65°C (5 min.)/  72ºC (7min.) | 65°C (5 min.)/  72ºC (7min.) | 65°C (4 min.)/  72°C (7 min.)/  72°C (5 min.) | 65°C (4 min.)/  72°C (5 min.) | 65°C (4 min.)/  72ºC (5min.)/ 72ºC (7min.)/  72ºC (3min.) |
| Cycles | 28-30 | 30-36 | 28-35 | 28-35 | 28-35 | 28-35 | 30-36 | 35 | 30-40 |

* 1 min with a ramp of 0-3 °C ^s–1^; ** Cycles (Denaturation, Annealing, Extension)
