## Supplementary figures and images for "Towards a species-level phylogeny for Neotropical Myrtaceae: notes on topology and resources for future studies"

### Supplementary Information 4

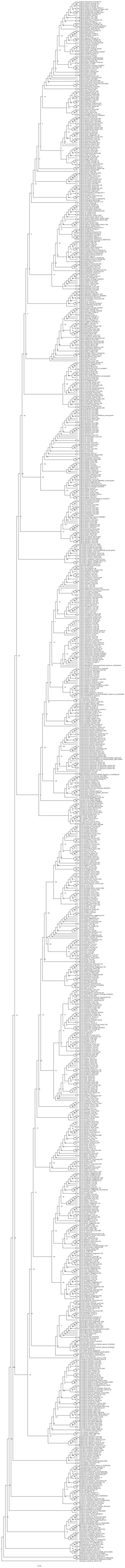
